# Clinical manifestations and phenotypic analysis of new ANK1 gene mutations in 12 Chinese children with hereditary spherocytosis

**DOI:** 10.1101/637553

**Authors:** Fei Xie, Lei Lei, Bin Cai, Lu Gan, Yu Gao, Xiaoying Liu, Lin Zhou, Jinjin Jiang

## Abstract

**Objective:** To summarize the clinical features and laboratory examination of ANK1 gene in 12 children with hereditary spherocytosis in China, and to determine the genetic mutations in those children.

**Methods:** The clinical data of children and their parents were collected and analyzed. The sequence of related genes was analyzed by second-generation sequencing technology. The suspected pathogenic mutations were detected by Sanger sequencing

**Results:** New mutations in the coding region of ANK1 was detected in 12 patients, which caused amino acid changes in the gene encoding, causing structural changes or loss of function.

**Conclusion:** ANK1 (c.1914_c.1918delTTTG), ANK1 (c.399T>G), ANK1 (c.1564delC), ANK1 (c.4439dupA<br>), ANK1 (c.4510_4513del), ANK1 (c.2961delC), ANK1 (c.2142dupT), ANK1 (c.2858+1G>C), ANK1 (c.3235delG), ANK1 (c.4739A>G), ANK1 (c.2638-2 A>G), ANK1 (c. 4739A>G) mutations in the coding region of the gene are the cause of suspicious disease in these 12 Chinese children. At the same time, second-generation gene sequencing technology is an effective means of confirming HS. Different types of mutations (P=0.388)and different mutation areas (P=0.660)had no significant effect on the severity of anemia. The 12 gene mutation sites in this study have not been included in the human genome database, dbSNP (v138) and ExAC databases. The new ANK1 gene mutations found in HS children can provide further exploration of the genetic etiology of HS in Chinese population.

## INTRODUCTION

Hereditary Spherocytosis (HS) is a kind of hemolytic anemia with erythrocyte membrane protein structure changes mainly characterized by jaundice, hemolysis and splenomegaly (1,2). It is also called congenital hemolytic anemia. The disease can occur in all parts of the world, and the incidence rate varies in different regions. The incidence of HS in Northern Europe is about 1/2000(3), while in China, the incidence of HS in adult is about 1/100,000 (4). The clinical manifestations of the patients varied greatly. The mild cases had no clinical symptoms and the severe cases could be manifested severe anemia, jaundice, splenomegaly, gallstones(5). For some patients with severe HS, repeated blood transfusions are often needed to maintain normal life and activities(6). And this brings a serious burden to families and society.

It is currently believed that HS-related mutant genes mainly include ANK1, SPTB, SPTA1, EPB42 and SLC4A1, and among these all related genes, ANK1 gene mutation is the most common(7). In terms of heredity, HS is mainly autosomal dominant, but there are also some reports of participatory recessive inheritance(8).

With the rapid development of post-gene sequencing technology and its application in medical detection, clinical gene sequencing technology has gradually shifted from first-generation sequencing to second-generation gene sequencing(9). In this study, 271 anemia-related loci were detected in 12 Chinese HS children by second-generation gene sequencing technology, and the detected new mutation sites were further confirmed using a generation of sequencing technology. This also proves the effectiveness, accuracy and convenience of second-generation gene sequencing technology in HS detection.

## Objectives and methods

### Patients

A retrospective analysis of patients from the First Affiliated Hospital of the Naval Military Medical University from May 2016 to December 2018, with a clear diagnosis of 12 children with HS based on clinical manifestations, hematology, and family history. All patients underwent second-generation sequencing, and children with detectable and predicted erythrocyte membrane protein gene mutations were included in the study. The diagnosis of HS is based on the literature standard(10). All children were included in different families and were not related.

## Methods

### Second-generation sequencing

2 ml of venous blood samples were taken from the children, and the whole genome DNA of the patients was extracted using DNA extraction kit (product of Beijing Tiangen Biochemical Technology Co., Ltd.) and using BloodGen Midi Kit (CWBIO, China). The DNA library was established according to the instructions of GenCap Custom Kit (product of Beijing MyGenostics Co., Ltd.), and 271 genes related to congenital anemia (including five erythrocyte membrane protein-coding genes related to the pathogenesis of HS) were selected by gene capture strategy. The biotin-labeled capture probe (80-120-mer) was used to cover all exons with non-repetitive regions. The average gene coverage of the targeted region was 94.86%, and the average sequencing depth was 436.75×,90.14% of targeted area coverage >30×, 60.29% of targeted area coverage >200×. The enriched DNA library was sequenced using the Illumina HiSeq sequencing platform. The sequencing results were compared with The Single-Nucleotide Polymorphism database (http://www.ncbi.nlm.nih.gov/projects/SNP/), the 1000 Human Genome Project (www.1000genomes.org), Exome Aggregation consortium (http://exac.broadinstitute.org/).

### Sanger sequencing

Sanger sequencing was performed on HS-related gene mutations detected by second-generation sequencing, compared to sites in the 1000 Genomes and ExAc databases with frequencies below 0.01 and functionally predicted to be detrimental. The genomic DNA of the patient’s parents was obtained for Sanger sequencing. Specific PCR amplification and purification of the PCR product was performed on the sample DNA, and the product was sequenced on an ABI 3730XL sequencer using a terminator cycle sequencing method. Variant sites were identified by comparing the DNA sequences to the corresponding GenBank (www.ncbi.nlm.nih.gov) reference sequences. The gene reference sequence transcripts are NM_001142446 (ANK1), NM_001024858 (SPTB), NM_003126 (SPTA1), NM_000342 (SLC4A1), and NM_000119 (EPB42), respectively.

### Pathogenicity prediction of mutation sites and analysis of protein function effects

application of Protein Variation Effect Analyzer (http://provean.jcvi.org/index.php), Sorting Intolerant From Tolerant (http://sift.jcvi.org), PolyPhen-2 (http://genetics.bwh.harvard.edu/pph2) and Mutationtaster (http://Mutationtaster.org) predict and analyze protein and structural hazards.

### Statistical methods

Statistical analysis was performed using SPSS 22.0. Non-normal measurement data is described in terms of median (range) and count data is described as a percentage. Comparisons between groups were performed using the Kruskal-Wallis H test. P < 0.05 was considered statistically significant.

## Results

### Clinical characteristics and laboratory results of children with HS in China

The clinical manifestations and laboratory results of 12 children with HS in China are shown in table 1. Among all children, 6cases were male (50%) and 6 cases were female (50%), and all of them were unrelated.11 cases (11/12,92%) were infected within 1 year of age. But the median age at which the disease of HS was diagnosed was 4 years and 8 months. In routine laboratory tests, these patients had a median reticulocyte percentage of 14% and a median total bilirubin level of 59.1 umol/L. According to the hemoglobin level at the time of treatment in our hospital with no blood transfusion for more than 30 days, the children with mild, moderate, and severe anemia were 2/12 (17%), 4 /12(33%), and 6 (50%).8 cases (8/12,67%) had a history of neonatal jaundice, however, there was no correlation with the severity of anemia in these children in the future(P=0.793). 10 cases (10/12,83%) had a history of blood transfusion; and splenomegaly was found in 7cases (7/12,58%).

**Table 1:**
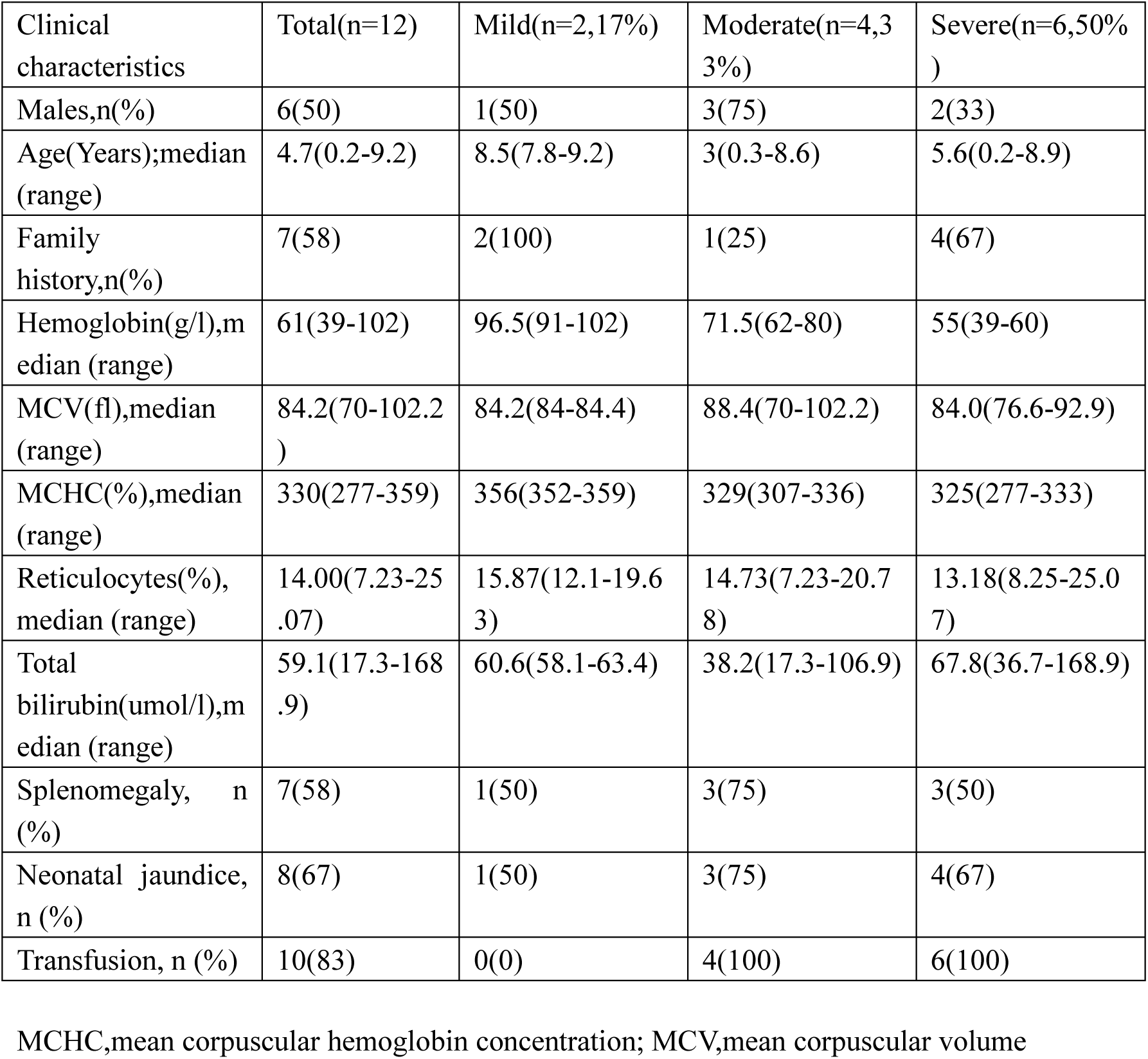
Clinical manifestations and routine laboratory findings of 12 children with newly diagnosed ANK1 mutations in China

| Clinical characteristics | Total(n=12) | Mild(n=2,17%) | Moderate(n=4,33%) | Severe(n=6,50%) |
| --- | --- | --- | --- | --- |
| Males,n(%) | 6(50) | 1(50) | 3(75) | 2(33) |
| Age(Years);median (range) | 4.7(0.2-9.2) | 8.5(7.8-9.2) | 3(0.3-8.6) | 5.6(0.2-8.9) |
| Family history,n(%) | 7(58) | 2(100) | 1(25) | 4(67) |
| Hemoglobin(g/l),median (range) | 61(39-102) | 96.5(91-102) | 71.5(62-80) | 55(39-60) |
| MCV(fl),median (range) | 84.2(70-102.2) | 84.2(84-84.4) | 88.4(70-102.2) | 84.0(76.6-92.9) |
| MCHC(%),median (range) | 330(277-359) | 356(352-359) | 329(307-336) | 325(277-333) |
| Reticulocytes(%), median (range) | 14.00(7.23-25.07) | 15.87(12.1-19.63) | 14.73(7.23-20.78) | 13.18(8.25-25.07) |
| Total bilirubin(umol/l),median (range) | 59.1(17.3-168.9) | 60.6(58.1-63.4) | 38.2(17.3-106.9) | 67.8(36.7-168.9) |
| Splenomegaly, n (%) | 7(58) | 1(50) | 3(75) | 3(50) |
| Neonatal jaundice, n (%) | 8(67) | 1(50) | 3(75) | 4(67) |
| Transfusion, n (%) | 10(83) | 0(0) | 4(100) | 6(100) |

MCHC,mean corpuscular hemoglobin concentration; MCV,mean corpuscular volume

### Genetic mutation characteristics of children with HS in China

The mutation characteristics of 12 newly diagnosed ANK1 Chinese children are shown in Table 2. Among the 12 children with new mutations, 6 cases (50%) were male and 6 cases (50%) were female, and there was no significant gender difference in the incidence of the disease. There were 7 cases (58%) with family history, 4 (57%) mutations from mothers, 3 (43) mutations from fathers, and all parents have a history of anemia. The remaining 5 patients (42%) were newly diagnosed with no family history. Among all the mutation types, the frameshift mutation, the shear mutation, the nonsense mutation and the missense mutation were 7 cases, 2 cases, 2 cases and 1 case, respectively. None of the mutations in all children were included in the genomic database, dbSNP (v138) and ExAC databases, nor in the literature.

**Table 2:** Gene mutation spectrum of 12 children with newly diagnosed ANK1 mutation in China

| Patients | Sex | age | Inheritance | Gene | Location | cDNA change | Protein change | Type |
| --- | --- | --- | --- | --- | --- | --- | --- | --- |
| 1 | male | 5 years and 3 months | De novo | ANK1 | Exon4 | c.399Thr>Gly | p.Tyr133X,1765 | Nonsense |
| 2 | female | 2 months | Hereditary | ANK1 | Exon17 | c.1914_1918delTTTGC | p.Pro638Profs*14 | Frameshift |
| 3 | male | 8 months | De novo | ANK1 | Exon14 | c.1564delC | p.Leu522Cysfs*44 | Frameshift |
| 4 | female | 3 months | Hereditary | ANK1 | Exon37 | c.4439dupA | p.Asn1480Kfs*20 | Frameshift |
| 5 | male | 9 years and 2 months | Hereditary | ANK1 | Exon37 | c.4510_4513del | p.Asn1504Trpfs*17 | Frameshift |
| 6 | female | 7 years and 11 months | Hereditary | ANK1 | Exon27 | c.2961delC | p.T988Pfs*30 | Frameshift |
| 7 | female | 8years and 11 months | Hereditary | ANK1 | Exon18 | c.2142dupT | p.Pro715Serfs*111 | Frameshift |
| 8 | male | 8 years and 7 months | Hereditary | ANK1 | Intron26 | c.2858+1G>C | - | Splicing error |
| 9 | female | 4 months | De novo | ANK1 | Exon28 | c.3235delG | p.Glu1079fs | Frameshift |
| 10 | male | 4 years and 1 month | Hereditary | ANK1 | Exon39 | c.4739A>G | p.Gln1580Arg | Missense |
| 11 | female | 7 years | De novo | ANK1 | Exon25 | c.2638-2 A>G | - | Splicing error |
| 12 | male | 3 months | De novo | ANK1 | Exon27 | c.2926C>T | p.Arg976Ter | Nonsense |

De novo, no family history with parents' genetic study

### Comparison of clinical symptoms in children with different membrane protein gene mutation types

The clinical symptoms of children with different membrane protein gene mutation types are shown in Table 3. Among the 12 new Chinese ANK1 mutations, the most common frameshift mutations were 7 cases (58%), followed by 2 cases of shear mutations (17%), 2 cases of nonsense mutations (17%) and one case of missense mutation (8%). In the degree of anemia corresponding to different mutation types, the median hemoglobin concentration was severe anemia in children with missense mutation, and the median hemoglobin concentration was moderately anemia in both the frameshift mutation, the shear mutation and the nonsense mutation, but this may be related to the less of cases in this study.

**Table 3:** Comparison of clinical symptoms in children with different membrane protein gene mutation types

| Clinical characteristics | Total(n=12) | Frameshift(n=7,58%) | Splicing error(n=2,17%) | Missense(n=1,8%) | Nonsense(n=2,17%) |
| --- | --- | --- | --- | --- | --- |
| Males,n(%) | 6(50) | 2(29) | 1(50) | 1(100) | 2(100) |
| Age(Years);median (range) | 4.7(0.2-9.2) | 0.7(0.2-9.2) | 7.8(7.0-8.6) | 4.1(4.1) | 2.8(0.3-5.3) |
| Family history,n(%) | 7(58) | 5(71) | 1(50) | 1(100) | 0(0) |
| Hemoglobin(g/l),median (range) | 61(39-102) | 62(53-102) | 64(56-71) | 39(39) | 62(44-80) |
| MCV(fl),median (range) | 84.2(70-102.2) | 84.0(70-86.5) | 91.6(90.2-92.9) | 76.6(76.6) | 92.5(82.8-102.2) |
| MCHC(%),median (range) | 330(277-359) | 333(323-359) | 307(306-307) | 277(277) | 338(324-351) |
| Reticulocytes(%),median (range) | 14.00(7.23-25.07) | 10.49(7.23-19.36) | 22.15(19.23-25.07) | 15.9(15.9) | 14.46(8.25-20.7) |
| Total bilirubin(umol/l),median (range) | 59.1(17.3-168.9) | 48.0(28.3-63.4) | 46.5(17.3-75.6) | 168.9(168.9) | 109.4(106.9-111.8) |
| Splenomegaly, n (%) | 7(58) | 3(43) | 2(100) | 1(100) | 1(50) |
| Neonatal jaundice, n (%) | 8(67) | 3(43) | 2(100) | 1(100) | 2(100) |
| Transfusion, n (%) | 10(83) | 5(71) | 2(100) | 1(100) | 2(100) |

MCHC,mean corpuscular hemoglobin concentration; MCV,mean corpuscular volume

### Distribution characteristics of new mutations in 12 Chinese children in the anchorage domain of ANK1

The distribution of 12 new ANK1 Chinese children in the anchoring domain of ANK1 is shown in Figure 1. Among them, 4 new mutations were located in the N-terminal domain, 5 were located in the central domain, and 3 were located in the C-terminal regulatory region. According to the degree of anemia, 4 cases (100%), 4 cases (50%), and 2 cases (67%) were severe anemia in the N-terminal domain, central domain, and C-terminal regulatory region. There was no significant difference in the severity of anemia of all new mutations in different regions(P=0.660).

**Figure 1:**
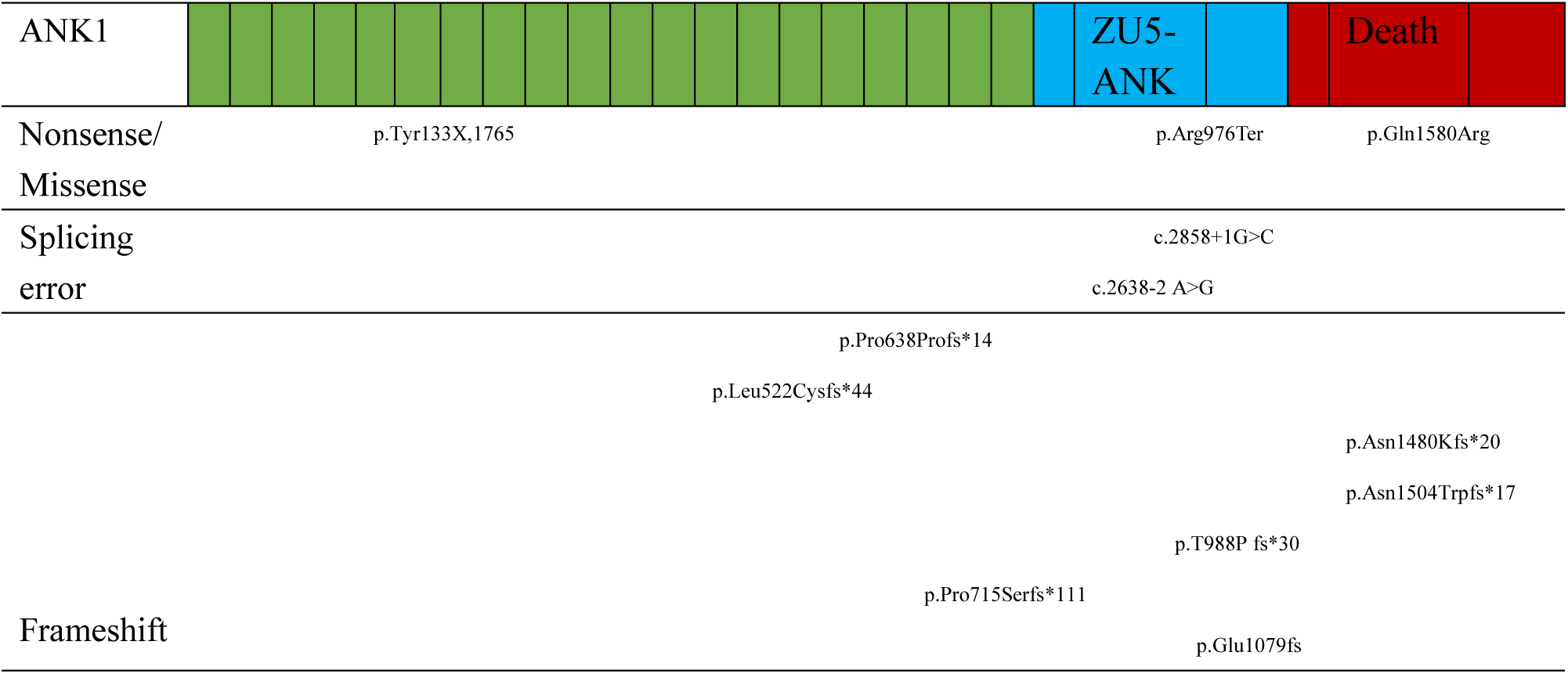
Schematic diagram of ANK1 anchoring domain

In the table, the green region indicates the N-terminal domain, blue indicates the central domain, and red indicates the C-terminal regulatory region.

## Discussion

Hereditary Spherocytosis (HS) is a kind of hemolytic anemia with erythrocyte membrane protein structure changes characterized by jaundice, hemolysis, hepatosplenomegaly and gallstones. In domestic relevant statistics, it accounts for about 84% of erythrocyte membranous diseases(11). In genetic terms, about 70% of HS patients have autosomal dominant inheritance, and autosomal recessive or new mutations account for about 25%(12–14). At the age of onset, HS can occur in any age group, but mostly in the age of neonatal or childhood, the median age of onset is about 5.6 years(15), and the incidence has no significant gender difference(16). In this study, 12 patients developed onset in neonates or infancy, suggesting a premature age of HS in children. The median age of all children is 4.8 years, which is consistent with previous studies. Because the clinical phenotypes determined by different pathogenicity loci are different, the clinical manifestations of HS in different periods are also different. In the fetal period, the phenotype of HS patients can be from no clinical symptoms to fetal hydrocephalus or even stillbirth. In childhood, HS often manifested as anemia, jaundice, hepatosplenomegaly, and most accompanied by a positive family history(17). Of the 12 patients in the study, 7cases (58%) had a parent with the same mutation gene locus and clinical manifestations of anemia, also meet the characteristics of HS having a positive family history in children. At the same time, the disease also has a high rate of misdiagnosis and missed diagnosis among children. Literature reports that the rate of misdiagnosis and missed diagnosis in children is 69.2% in China (18). Therefore, there are still many shortcomings understanding and diagnosis of the disease in China. In this study, in all children with splenomegaly, the median age of onset was 7 years old, Infancy of HS often had no clinical symptoms of splenomegaly. All patients had no gallstone formation during the examination, which was different from the clinical manifestations of adult HS patients, which may be related to the long-term chronic process of splenomegaly and gallstone formation. For children with younger age, because of the lack of characteristic clinical manifestations, this makes clinical diagnosis difficulty, increases the rate of misdiagnosis and missed diagnosis, and also delays treatment. Therefore, for children with early onset and short duration, laboratory tests and genetic tests are necessary for the diagnosis and identification of the cause.

The research indicates that the sites related to HS gene mutation are mainly located in the following genes: ANKl, SPTB, SPTA1, EPB42, and SLC4A1. Among them, the most common type of mutation is ANK1 mutation, which accounts for about 50% of all HS gene mutation types(19–21), followed by SLC4Al and SPTB locus mutations(22,23). There are few related studies on HS gene in China, Most studies are case reports and lack of large sample studies (24–26). It is also rare research and large-scale studies on children’s ANK1 gene in other countries (7). The 12 ANK1 mutation found in this study are all mutations unreported. All gene mutation sites have not been included in the thousands of human genome database, dbSNP (v138) and ExAC databases, and have not been reported in the literature. There is no report of new gene ANK1 mutation similar to our study in China’s children’s with HS, which also suggests that ANK1 gene mutations are less studied in Chinese HS. And the type of ANK1 mutation in Chinese HS children has the uniqueness of its mutation type compared with other regions and other ethnic groups, whcih deserve further study.

In terms of structure, the corresponding altered of erythrocyte membrane structural proteins are also different for different mutant genes. The most common erythrocyte membrane structural proteins are ankyrin 1 (ANK1 encoding) and spectrin β-1 (SPTB gene encoding), spectrin alpha-1 (SPTA1 gene encoding), erythrocyte protein 4.2 (EPB42 gene encoding) and solute carrier family 4-1 (SLC4A1 gene encoding)(8,27). For the ankyrin encoded by ANK1, it is mainly expressed in the erythrocyte membrane by linking an anion channel, a Rh complex and a contracted ovalbumin(27). In the erythrocyte membrane structure, ankyrin binds to the self-joining point of the β-chain tail of the contractile protein at one end, and the other end is linked to the band 3 protein, which fixes the membrane skeleton in the lipid bilayer and plays an important role in stabilizing the erythrocyte membrane(28). Ankyrin typically consists of three domains: an N-terminal domain containing multiple ankyrin repeats, the central region containing a spectrin binding domain and a C-terminal regulatory domain. Studies have shown that patients with ANK1 mutations in the spectrin binding domain have the most severe anemia compared to mutations in other domains(7). Among the 12 children with ANK1 mutation reported in this article, 5 patients had a mutated region in the central region, and 4 of them had moderate to severe anemia. This indicates the severity of anemia after mutation in this region. However, in this study, children with moderate to severe anemia accounted for 100% and 67% of all mutations in the N-terminal domain and the C-terminal regulatory region respectively. This result is different from the more severe anemia in patients with mutations in the central region of previous studies, which may be related to the specificity and regionality of Chinese children’s races, and further research is needed for confirmation. Studies have shown that the higher the affinity of the ankyrin-encoded ankyrin to beta-spectrin, the higher the shape and stability of the erythrocyte membrane(29). Conversely, when the quantity or quality of of ankyrin is reduced or deficient, its role in stabilizing the cell membrane lipid bilayer decreases, loss of part of the lipid, causing a decrease in the area of the red blood cell membrane, resulting in a change in the shape of the red blood cells(30). Therefore, when ANK1 is mutated, its encoded ankyrin structure also changes, which reduces the plasticity and stability of the erythrocyte membrane, thereby accelerating the dissolution and destruction of red blood cells.

There are more than 60 kinds of human HS-related ANK1 gene mutations reported(31). The 12 ANK1 gene mutation sites reported in this paper are located in ANK1 (c.1914_c.1918delTTTG), ANK1 (c.399T>G), ANK1 (c.1564delC), ANK1 (c.4439dupA <br>), ANK1(c.4510_4513del), ANK1(c.2961delC), ANK1 (c.2142dupT), ANK1 (c.2858+1G>C), ANK1 (c.3235delG), ANK1 (c.4739A>G), ANK1(c.2638-2 A>G) and ANK1(c.2926C>T), The above-mentioned mutation sites were also confirmed by Sanger sequencing, and the Provean software predicted that the protein structure was harmful. In subsequent gene analysis of the family we found that in ANK1 (c.1914_c.1918delTTTG), ANK1 (c.4439dupA<br>), ANK1 (c.4510_4513del), ANK1 (c.2961delC), ANK1 (c.2142dupT), ANK1 (c.2858+1G>C) and ANK1 (c.4739A>G) mutations, the father or mother of the child carries the same genetic mutation site as the child, and also has a history of anemia. These mutation sites are in accordance with the genetic pattern of HS autosomal dominant inheritance, so it can be considered that the above new mutation sites are the pathogenic gene mutation sites of these children. In the families of children with ANK1 (c.399T>G), ANK1 (c.1564delC), ANK1 (c.3235delG), ANK1 (c.2638-2 A>G) and ANK1 (c.2926C>T) mutations, the parents of the children were all normal wild-type genes and had no history of family anemia. Combined with the clinical manifestations and laboratory findings, the mutation sites of these children were considered to be new pathogenic mutations. In all children with new mutations, frameshift mutation, shear mutation, nonsense mutation, and missense mutation were 7 cases (58%), 2 cases (17%), 1 case (8%), and 2 cases (17%). Among the 7 children with frameshift mutation, 4 (57%) children with moderate to severe anemia, 2 children withSplicing errorwere moderate to severe anemia, 1 child with missense mutation was severe anemia, and 2 children with nonsense mutation were moderate to severe anemia. This may be related to the complex condition of most children who come to our hospital, most of the children who had visited several hospitals for multiple times and had not yet been able to identify the cause. Because of the small number of different types of mutations mentioned above, it is unable to explain the relationship between different types of mutations and the severity of anemia. Therefore, follow-up case collection is still needed to further confirm the correlation between different types of mutations and the degree of anemia. The above 12 children with genetic mutation sites have not been included in the 1000 human genome database, dbSNP (v138) and ExAC databases, and have not been reported in the literature. Therefore, we can regard that the ANK1 mutations in the above 12 children are all new mutation sites in Chinese children’s HS.

The application of gene detection technology in various hereditary diseases is gradually becoming popular. It has more precision and its role can not be replaced by other laboratory tests for some hereditary diseases which are difficult to diagnose (32,33), therefore, it has been gradually applied to the detection of hereditary erythrocyte membrane diseases, including HS(34). The second-generation gene sequencing technology provides an economical, efficient, rapid, and direct method for detecting complex and diverse genes, especially for patients whose routine laboratory tests cannot identify the cause or multiple blood transfusions(35). The second-generation gene sequencing technology provides a more accurate and rapid detection method for the identification of its cause. The results of this study suggest that this technique is of great significance in the diagnosis of HS. It is a powerful means of distinguishing between congenital hemolytic anemia and immune anemia caused by other causes, especially for children with new family mutations who have no family history, which can help diagnose early, clear the condition and relieve parents’ anxiety, choose the right treatment according to the severity of the disease.

